# Designing synthetic bacterial consortia for landfill leachate treatment based on community matrices and regression tree analysis

**DOI:** 10.1101/543694

**Authors:** Georgeos Hardo, Esther Karunakaran, Narciso A. Couto, Andrew P. Beckerman, Jagroop Pandhal

**Author notes:** Correspondence: Jagroop Pandhal, Department of Chemical and Biological Engineering, University of Sheffield, Mappin Street, Sheffield S1 3JD, United Kingdom.

## Abstract

The performance of microbial communities exploited by industry are largely optimised by manipulating process parameters, such as flow rates, growth conditions, and reactor parameters. Conversely, the composition of microorganisms used are often viewed as a “black box”. This is mostly due to the relatively high costs and technical expertise required to identify and quantify the microbial consortia, as well as limited tools to create functional assemblages. Unknown details about the interactions among species may impose a limit on how much microbial function can be optimised for industrial purposes. Here, a new workflow was developed for studying microbial consortia using high throughput, species and community specific measurements of growth rates and yields. Growth rate and yield among all single, pairwise, triple, quadruples, quintuple and sextuple combinations of six bacterial isolates on landfill leachate were evaluated. Additive, antagonistic (e.g. competitive) or synergistic (+/-) interactions can be inferred from the rate and yield data. We found that antagonistic interactions, which hinder growth and yield, were the dominant interaction type, with only a few synergistic interactions observed. Mixed effects models were used to investigate the relationship between interaction type and species richness (biodiversity). Community identity was found to be a more important factor in predicting yield determining interactions but not rate determining interactions. Species richness was a good predictor of rate determining interactions, with the most positive interactions happening at a low species richness. Regression tree analysis identified Lysinibacillus sp. as a keystone species, a genus previously associated with bioremediation. Its presence led to a drastic change in the function of the synthetic ecosystem, with both positive yield and rate determining interactions. We were able to infer interactions about specific pairs of species, and the competitive/synergistic tendencies of single species from only basic top-down growth measurements. In this way, we have demonstrated how factorial experiments using isolated microorganisms can be used to ultimately design synthetic consortia with desirable traits for industry.

**Funding statement:** This research was funded by the EPSRC (Vacation Bursary Award), BBSRC (Impact Acceleration Award), Department of Chemical and Biological Engineering, The University of Sheffield (Summer Undergraduate Research Fellowship) and Viridor Ltd.

**Ethics statements:** (Authors are required to state the ethical considerations of their study in the manuscript, including for cases where the study was exempt from ethical approval procedures)

*Does the study presented in the manuscript involve human or animal subjects:* No

**Data availability statement:** Generated Statement: This manuscript contains previously unpublished data. The name of the repository and accession number are not available.

## 1 Introduction

Industrial bioprocessing using microorganisms often use single species or strains and is most frequently optimised through either strain engineering or process manipulations. Improvements in process efficiency are achieved through process parameter optimisation (Simutis and Lübbert 2015) or culture media optimisations (Singh et al. 2017). Strains can be modified using either traditional genetic and metabolic engineering, or through emerging data driven synthetic biology approaches (Peralta-Yahya et al. 2012). However, single species bioprocesses are inherently limited by the metabolism – engineered or not – of the strain used (Pandhal and Noirel 2014). Microbial communities found in nature are less restricted in their metabolic abilities, and can often undertake metabolically complex tasks. These synergies are driven by the ability of individual species to share by-products between one another (Chiu, Levy, and Borenstein 2014). Microbial communities are also more robust compared to single species cultures as they demonstrate emergent properties, reflecting the consortium’s ability to adapt to changing process conditions or the presence of diverse metabolites.

Highly complex microbial communities are routinely harnessed for industrial processes, particularly for wastewater treatment or for biogas production. Complex consortia with diverse metabolic capabilities and functional characteristics are extremely valuable in these contexts, where feedstocks (i.e. waste) can vary or environmental parameters of the process are difficult to control. Microbial consortia with desirable functions are often selected by enrichment by providing suitable industrial conditions, or even by seeding from similar natural environments. However, rectification of process failures are inherently empirical and limited to changing process conditions because the tools required for fully characterising the interactions between the microorganisms that make up the consortia remain limited. Optimisation of the community of microbes can be undertaken by adaptive evolution, if a suitable selective pressure can be identified, or by construction of synthetic consortia from laboratory isolates. However, construction of synthetic consortia may present difficulties due to the presence of so-called “unculturable bacteria” (Stewart 2012). These are bacteria that are difficult to grow in a laboratory, either due to lack of knowledge of their metabolic requirements or their reliance on other community members. With increasing knowledge of the metabolic requirements and the availability of techniques to culture these difficult organisms, laboratory-based methods to isolate and then create synthetic consortia to outperform existing microbial communities is an incipient approach to optimise industrial processes and enable additional functionality, for example, degrading new manmade materials or recovery of resources from waste streams.

Engineered synthetic consortia should seek to mimic or outperform the function of the wild type consortium. A key objective may be to find the smallest number of species which meet bioprocessing requirements. The minimal consortium would contain only the metabolic functioning and diversity required to meet the end goal of the process. An efficient consortium could entail individual species which do not compete for resources in such a way that the overall growth of the community is slowed or hindered, and it is for this reason that natural consortia are often not ideal. During laboratory investigations to identify appropriate members of a synthetic consortia, it would be desirable to find combinations of bacteria which will naturally interact synergistically or have metabolic requirements which complement one another. Synergistic interactions between two or more different strains may be inferred through an increase in growth rate over and above the sum of the individual growth rates or in an increase in overall yield, over and above the sum of the individual yields.

In this study, we aimed to identify bacterial species that can be utilised to construct synthetic microbial consortia capable of effective bioremediation of landfill leachate. Leachate is generated when liquid (most often rainwater) soaks through the landfill, solubilising compounds which diffuse from the waste into the water including heavy metals such as lead and mercury, organic matter in solution (alcohols, aldehydes), sulphates, chlorides, ammonia including other inorganic components, and many xenobiotic compounds (such as halogenated organic compounds) (Kjeldsen et al. 2002). We isolated bacteria from a man-made landfill leachate habitat and examined the effects of competition and synergistic interactions in synthetic combinations of six bacterial isolates via in depth analyses of yield and growth rates. We identified consortia able to grow at a faster rate or reach a final higher biomass yield than that of the original community, using landfill leachate as the only carbon source. We explored the effects of strain identity on performance using regression tree analysis to identify key bacterial players in the synthetic landfill leachate ecosystems constructed. This novel data analysis approach, combining regression tree analysis and mixed models reveals the effects of individual species and species richness on the degradation rate and yield, and can be applied to other consortia in different environments.

## 2 Materials and Methods

### 2.1 Retrieving isolates from leachate

1 L of landfill leachate was collected from two locations at a local landfill site (Erin, Derbyshire, U.K.) on 27^th^ January 2016 in a sterile glass container. Upon receipt in the laboratory, the leachate was mixed well and 1 mL of the leachate from each location was spread on to R2A agar. The plates were incubated at 25°C for 48 hours. Thirteen colonies with a distinct colony morphology and appearance were transferred to fresh R2A agar. The plates were incubated at 25°C for 24 hours. A single colony from each of the thirteen plates were used to inoculate 5 mL of R2A broth. The cultures were allowed to incubate at 25°C for 24 hours. The isolates were collected by centrifugation at 4500 g for 5 minutes. The supernatant was discarded and the isolates were resuspended in 1 mL R2A broth supplemented with 10% (v/v) sterile glycerol. The suspension was stored in the −80°C freezer until further use.

### 2.2 DNA extraction

The 13 isolates were taken from the glycerol stocks and streaked onto R2A agar plates and incubated at 30°C overnight. A sterile pipette tip was used to transfer cells from the plates into 10mL of M9 salts supplemented with 20% w/v acetate.

The Cetyltrimethyl Ammonium Bromide (CTAB) protocol for DNA extraction was used to extract the DNA from each isolate according to the protocol as described previously (Karunakaran et al. 2016). DNA was quantified using a spectrophotometer (Thermo Scientific Nano Drop 2000, Loughborough, UK).

### 2.3 16S gene DNA sequencing

DNA from each isolate was taken for PCR amplification. Preliminarily, a low fidelity PCR was carried out to confirm that all isolates were bacterial. GoTaq Green Master Mix from Promega UK was used with 16S rDNA PCR primers in the initial run. The 27F (5’-AGAGTTTGATCMTGGCTCAG) and 1492R (5’-TACGGYTACCTTGTTACGACTT). A gel electrophoresis on 1.5% agarose with ethidium bromide was run with the amplified DNA at 130 volts for 45 minutes. A 1 kilobase molecular-weight size marker (Hyperladder 1KB from Bioline UK) was used to confirm that the 16S ribosomal RNA hypervariable region had been isolated (supplementary material). A high fidelity PCR was performed with the same 16S primers but using Q5 Hot start 2x master mix (New England Biolabs, Hitchin, U.K.). After amplification, an Applied Biosystems 3730 DNA Analyser using BigDye v3.1 was used to sequence the genes for the 16S hypervariable region. Identification was performed using the NCBI Nucleotide BLAST and searching the Nucleotide Collection (nr/nt) database. Both forward and reverse sequences were used in the search to confirm to the highest level of confidence the identification.

### 2.4 Growth measurements

A modification of the standard M9 salts protocol (Sambrook and W Russell 2001) was used to cultivate the cells, where glucose/acetate was replaced with landfill leachate as the only carbon source (hereon referred to as leachate growth media). A 5 × M9 minimal salts solution (Sigma-Aldrich, Dorset, U.K.) was prepared by dissolving 56.4 g of M9 into 1L of water of distilled water and autoclaved at 121°C for 15 minutes. 200 mL of 5X M9 salts were then transferred to a separate autoclaved 1L Duran bottle, and to it added 2mL of 1 M magnesium sulphate and 100 μL of 1 M calcium chloride, both filter sterilised using a 0.22 μm syringe filter. 100 mL of landfill leachate was centrifuged to remove particulate matter and sterilised using a 0.22 μm syringe filter. It was then added and made up to 1 L using autoclaved distilled water.

Bacterial pre-cultures were grown to a sufficiently high optical density (OD) of 0.6, at 37°C for 4-8 hours with shaking at 250 RPM. A 1 mL aliquot was used to measure OD of the sample in a spectrophotometer at 595 nm. A further 1 mL aliquot from was transferred to an autoclaved microcentrifuge tube and spun at 10,000 g for 5 minutes. The supernatant was removed and the pellet was washed by gently pipetting up and down with 1 mL of leachate growth media. The sample was then spun down once more at 10,000 g for 5 minutes. The supernatant was removed and then replaced once again with 1 mL of leachate growth media, pipetting gently up and down to re-suspend the cells. Each aliquot was diluted to 0.01 OD595. Aliquots from these were used to prepare the combinations of bacteria with a required starting OD of 0.01 (595 nm). For pairwise combinations 500 μL of each isolate at 0.01 OD595 were taken and mixed to make a final volume of 1 mL. For three-way combinations, 350 μL of each isolate were mixed for a final volume of just over 1 mL.

A TECAN GENios (Seestrasse, Switzerland) was used for the growth measurements. 29 combinations per plate were tested with negative controls (un-inoculated leachate growth media). The spectrophotometer was set to measure scattering at a wavelength of 595nm every 20 minutes for up to 72 hours, with orbital shaking for 1000 seconds before each measurement to ensure homogeneity in each well during the measurement.

The exponential growth rates were calculated using Growth Rates software (Hall et al. 2014), but also checked manually. The yield was defined as *OD_final_* – *OD_initial_*. The implicit assumption here is that growth can be used as a proxy for the breakdown of organic components, as these provided the sole source of carbon to the bacteria, and hence growth can be an indication of bioremediation.

### 2.5 Statistical analyses

Rate and yield factors for each consortium were calculated to analyse whether growth and yield were determined by antagonistic, additive or synergistic effects (Fiegna et al. 2015). The rate factor was defined by:

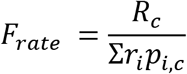

where *R_c_* is the observed growth rate of the consortium itself, and *r_i_* is the growth rate of isolate *i* in a monoculture, and *p_i,c_* is the presence of isolate *i* in consortium *c* (either 0 or 1). Similarly, the yield factor was defined by:

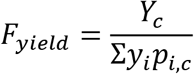

where Y is yield instead of growth rate.

We investigated the effect of consortia diversity on rate and yield using regression with random effects via the lme4 package for R (Bates et al. 2015). Our model fit rate or yield as a function of diversity (number of species in the consortia) with a random effect specifying the community composition (e.g. a consortia of three species is represented by 20 different combinations of the six pool species). *F_rate_* as a function of species richness was modelled as a power law and *F_yield_* as a function of species richness was modelled as a 2^nd^ order polynomial.

We explored the potential identity of keystone species in consortia using a regression tree analysis via the rpart package for R (Therneau et al. 2015). Using fitted values from the regression models of *Prate* and *F_yield_*, we recursively partitioned these data by diversity and species identity. This process identifies combinations of richness and species identity that are associated with maximal or minimal rate and yield.

## 3 Results

### 3.1 Isolation and identification

Successful PCR amplification with 16S primers revealed that all cultured isolates taken from the landfill leachate were prokaryotic. Of the 13 isolates taken from the landfill site, 9 were identified at least at the genus level and 6 were unique. Following NCBI Nucleotide BLAST searching using the Nucleotide Collection (nr/nt) database, three strains of *Pseudomonas* were found and were grouped into one OTU, hence the total number of identifications for this study was kept as 6 (Table 2). Inconclusive sequences were most likely not axenic isolates.

**Table 1.** List of isolates identified to at least the genus level. Isolates 8, 9 and 12 were part of the same genus. These were grouped into the same operational taxonomic unit (OTU).

| Isolate ID | Identification | Family | Phylum |
| --- | --- | --- | --- |
| F1 | *Brevundimonas diminutia* | Caulobacteraceae | Proteobacteria |
| F4 | *Lysinibacillus sp.* | Bacillaceae | Firmicutes |
| F7 | *Alcaligenes sp.* | Alcaligenaceae | Proteobacteria |
| F8 | *Pseudomonas sp.* | Pseudomonadaceae | Proteobacteria |
| F10 | *Paenibacillus sp.* | Paenibacillaceae | Firmicutes |
| F13 | *Stenotrophomonas chelatiphaga* | Xanthomonadaceae | Proteobacteria |

### 3.2 Are growth and yield additive, antagonistic, redundant or synergistic?

Figure 1 plots observed growth rates and yields vs. rates and yields expected if consortia were performing additively. The 1:1 line implies additive effects, and consortia which are placed above this line are performing with synergy and those below with antagonism (e.g. competitive; rates and yield are less than additive). Higher rates indicate faster toxic organic molecule breakdown, and higher yields indicate a higher carrying capacity of the landfill leachate (corresponding to a larger subset of the organic compounds being degraded).

**Figure 1.**
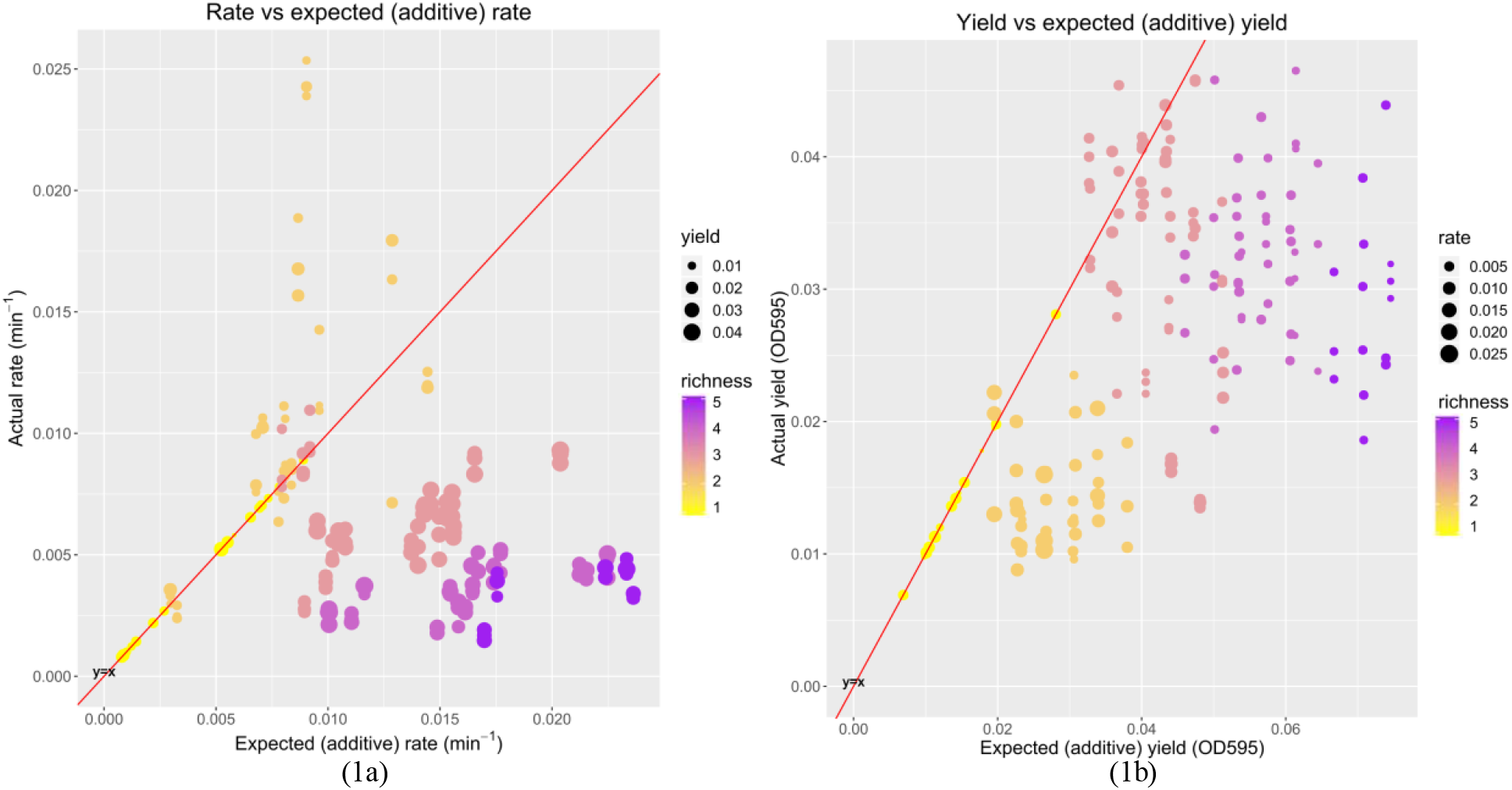
Observed (measured) versus expected (additive) growth parameters. Left: Shows the observed rate vs the expected (additive) rate of each consortium (each represented by a dot) with colour denoting species richness and dot size denoting yield. Right: Showing the observed yield vs the expected (additive) yield of each consortium (each represented by a dot) with colour showing species richness and dot size showing rate. The purpose of the colour and dot size is to capture all four dimensions of the data in a single image. The red 1:1 line indicates the threshold for inferring positive overall interactions. Consortia lying above the 1:1 line have an inferred interaction of ***F_rate_*** > **1** or ***F_yield_*** > **1**, indicating positive or synergistic interactions whilst consortia lying below the 1:1 line have an interaction of ***F_rate_*** <1 or ***F_yield_*** < 1, which indicates a negative interaction. Consortia on or near the line have interaction factors of ***F_rate_*** ≈ **1** or ***F_yield_*** ≈ **1**, which implies no overall interaction. For this reason, all monocultures (shown in yellow) lie on the 1:1 line as they have no interaction with any other species.

In Fig 1a, consortia with three or more species appear to be growing with less than additive rate performance, while the consortia of two species appeared to be growing with synergistic rate performance. 21% of consortia showed some positive rate determining interaction (*F_rate_* < 1). In Fig 1b, nearly all consortia were demonstrating less than additive (antagonistic) yield; only 9% of consortia showed positive yield determining interactions, most of which had a species richness of 3.

Positive or synergistic interactions were more likely to be found in 2-way microbial consortia, with 69.2% of 2-way microbial consortia having rate determining interaction factors (*F_rate_*) greater than 1. 10.5% of 3-way microbial consortia had *F_rate_* > 1, and 0% of both 4-way and 5-way microbial consortia had *F_rate_* > 1. Finding instances where yield determining interaction result in a positive interaction, *F_yield_* > 1, is less likely than for growth rate. 5.1% of 2-way consortia, and 21% of 3way consortia showed positive interactions. 0% of 4 and 5-way consortia showed positive interactions. Of all consortia, only one showed both positive rate and yield interactions, this was a 2-way consortium of *Lysinibacillus sp.* and *Alcaligenes sp*..

### 3.3 Are rate and yield related to consortia diversity?

The mean *F_rate_* and *F_yield_* values were plotted against species richness in Figure 2 where it is possible to see the negative power relationship between rate determining interactions and species richness and the arc-shaped relationship between yield determining interactions and richness. The results of our modelling are presented below.

**Figure 2.**
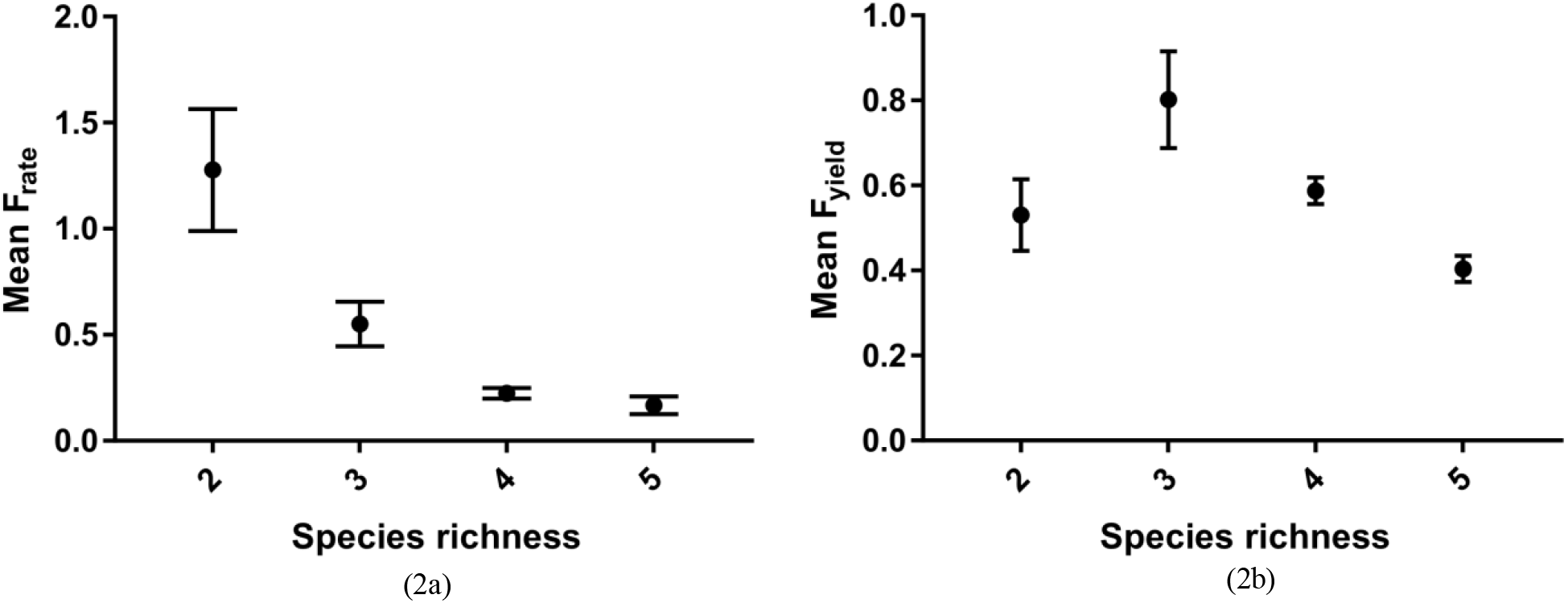
Left: Mean rate determining overall interaction factor vs species richness with error bars showing the 95% confidence intervals. A species richness of 1 is not plotted because by definition the interaction factor of a monoculture is always 1 – and it is meaningless to talk about inter-species interactions for a single species culture. Right: Mean yield determining overall interaction factor vs species richness with error bars showing the 95% confidence intervals. Species richness of 1 is not plotted again because the interaction factor would be 1. It should be noted that there are fewer 4-way combinations than 3 way combinations, and fewer 5-way combinations than 4-way combinations due to the finite way of choosing *k* elements from a set size *n*.

As detailed in the methods, we fit a power law model to the rate factor vs biodiversity data given by 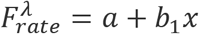 with λ=-0.25, a=0.5109, b_1_=0.228. Rate factor declined non-linearly with species richness (p<0.01, adjusted R^2^ = 0.82). We also fit a second order polynomial to the yield factor vs biodiversity data which was given by *F_yield_* = a + b_1__x + *b*_2_, with a=-0.581, b_1_=0.830, b_2_=-0.130 (p<0.01, adjusted R^2^ = 0.25). The random effect of consortium composition captured substantial variation compared to a model without the random effect and thus improved inference via our fixed effects (Δ*AIC_rate_* = −166,Δ*AlC_yield_* = −141).

### 3.3 Are there keystone species?

Our regression tree partitioning of the effects of diversity and species identity (Figures 3 and 4) revealed that the highest growth rate interactions were from low diversity (<3 species) consortia missing species 13, *Stenotrophomonas chelatiphaga.* Interestingly, the next highest growth rate factor was for consortia with <3 species containing *Stenotrophomonas chelatiphaga.* This reveals that the dominant “force” affecting consortium growth rates is derived from species richness. While it is seen that the presence of *Stenotrophomonas chelatiphaga* in 2-way consortia causes decreases in positive interactions, the effects of this change are damped by larger effect of species richness. This is what is expected given the modelling results for rate factor vs biodiversity.

**Figure 3.**
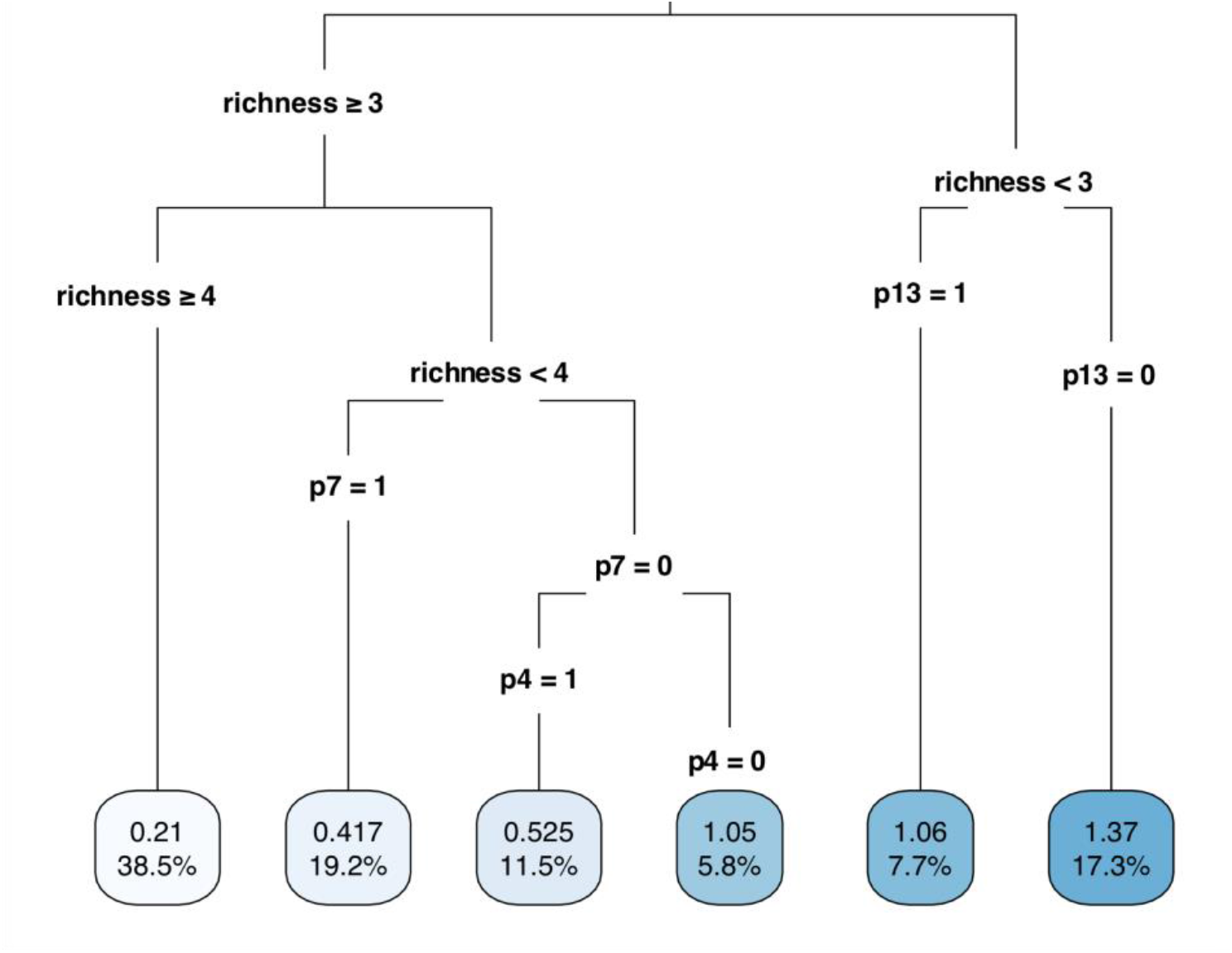
Rate factor regression tree made with the rpart package in R (Therneau et al. 2015). The leaves at the bottom are the rate factors and each partition shows the threshold where the data was split, and which variable. The partitions in this tree are by species richness and species presence. Presence is quantified by either 0 or 1 for each isolate, for example, p7=0 means *Alcaligenes sp.* is not present. The left of the tree contains the consortium data for consortia with a richness or 3 and above, whilst the right of the tree contains data for consortia with a species richness of 2. The data set used to train this regression tree does not contain the monoculture data because this would simply be a single redundant branch from the tree with richness=1 leading to a rate factor of 1. Refer to table 2 for isolate names.

In Figure 3, the data is initially split by richness. The right of the tree shows the 2-way consortia are further partitioned by the presence of *Stenotrophomonas chelatiphaga*. When *S. chelatiphaga* is present, an average rate determining interaction factor of 1.06 is observed, however when it is not paired with any other isolate the interaction factor rises to 1.37. The branch on the very right-hand side of the tree with richness >=4 is merely reiterating what is seen in Figure 2a, high species richness (4 and 5) leads to very low rate determining interaction factors, implying more competition. The branch for consortia with a species richness of 3 is then split based on the presence of isolate 7, *Alcaligenes sp.* whereby the presence of this species results in an overall lower mean *F_rate_*. The same is true for consortia which do not contain *Alcaligenes sp.* but do contain isolate 4, which is *Lysinibacillus sp.* It is implied that *Lysinibacillus sp.* is so competitive that when removed from 3-way consortia the average *F_rate_* increases from 0.525 to 1.05 on average.

Our regression tree partitioning of the effects of diversity and species identity on yield interactions revealed that the most positive yield interactions are associated with *Lysinibacillus sp* and *Paenibacillus sp* and diversities of 3 and 4 species. Figure 4 shows the regression tree for determining factors contributing to the yield factor. The tree is first split by the presence of isolate 4, *Lysinibacillus sp*.. The right shows that when *Lysinibacillus sp.* is present, the average interactions which determine yield are much more positive than when it is not present. Branching off from p4=0 there is another branch indicating the lack of isolate 7, *Alcaligenes sp*.. This branch has the second lowest yield factor, implying the second most competitive ecosystem for high yields. This occurs when both *Alcaligenes sp.* and *Lysinibacillus sp.* are not present in the consortia. When these two isolates are absent they lead to high implied synergistic interactions for growth rate but high implied competition for yield, and when they are present they lead to overall higher implied synergistic interactions for yield but overall much higher implied competition for rate. However, when these two isolates are alone they form the only consortium to have both positive rate and yield determining interactions. Much of the right-hand side of the regression tree in Figure 3 is devoted to showing the differences in yield determining interactions arising from changes in species richness where *Lysinibacillus sp.* is present in the consortium. When it is present, it is correlated with more synergistic interactions in consortia of species richness 2 and 3. Interactions are relatively synergistic when both it and isolate 10, which is *Paenbacillus sp,* are paired up with any other isolate to form a 3-way combination – resulting in an average yield factor of 1.07. This implies that *Paenbacillus sp* and *Lysinibacillus sp.* will either weakly positively interact or not interact with the third isolate in the system.

**Figure 4.**
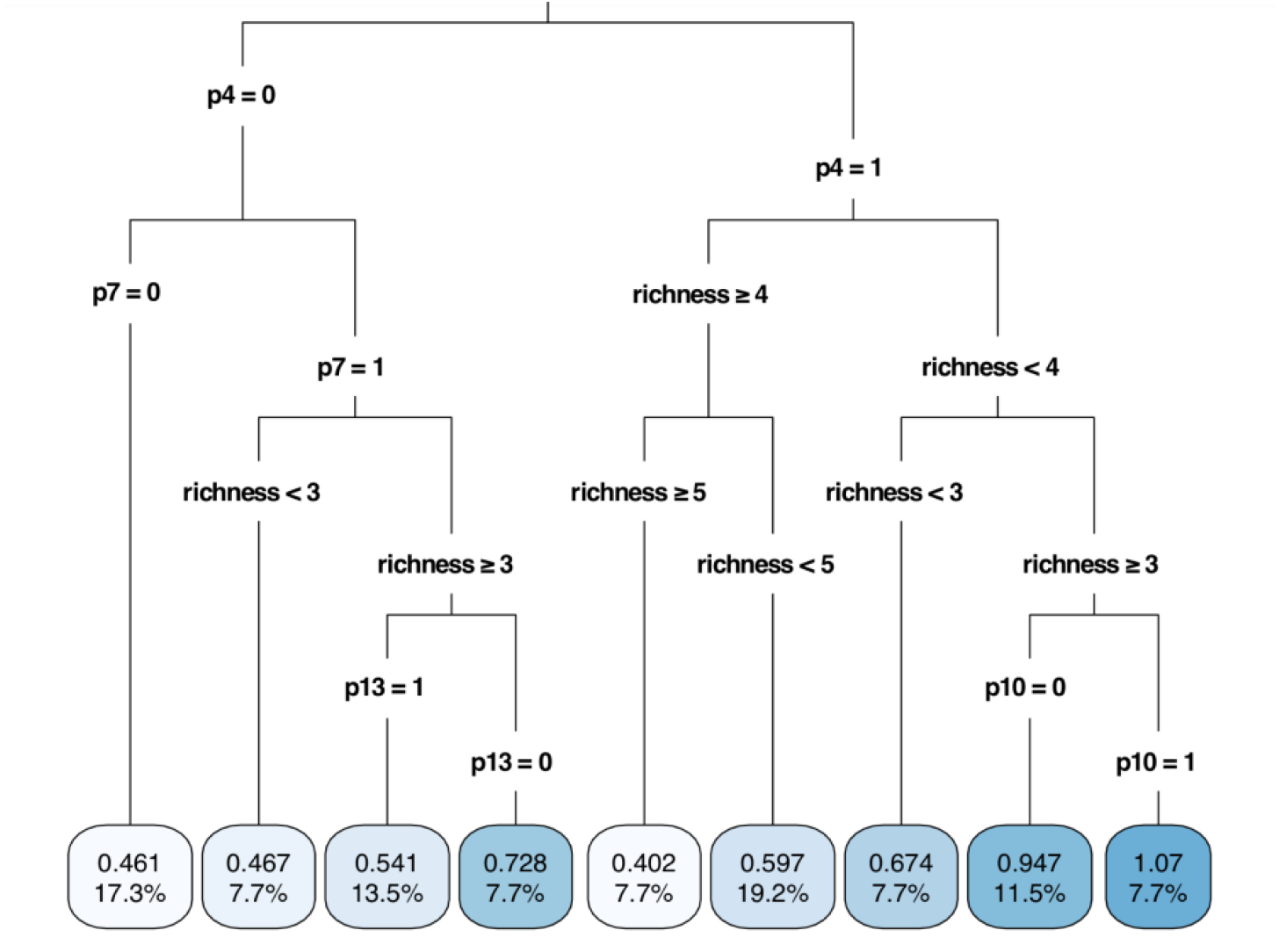
Regression tree made with the rpart package in R. The leaves at the bottom of the tree are the yield factors. The partitions in this tree are by species richness and species presence. Presence is quantified by either 0 or 1 for each isolate, for example, p7=0 means *Alcaligenes sp.* is not present. The left of the tree contains the consortium data for consortia with species 4 *(Lysinibacillus sp.)* not present and the right of the tree contains consortia where it is present. The data set used to train this regression tree does not contain the monoculture data because this would simply be a single redundant branch from the tree with richness=1 leading to a rate factor of 1. Refer to table 2 for isolate names.

In addition to these data, we further explored how rate and rate factor were correlated, and how yield and yield factor were correlated, to determine whether systems with high rates or yields also exhibit high synergistic interactions and vice versa. Linear models were fitted to the rate factor vs rate data, and it was found that positive interactions were correlated with growth rate (P < 0.001, R^2^=0.79). Yield was less strongly correlated with interaction positivity (P < 0.001, R^2^=0.46). These data are represented in Figure 5.

**Figure 5.**
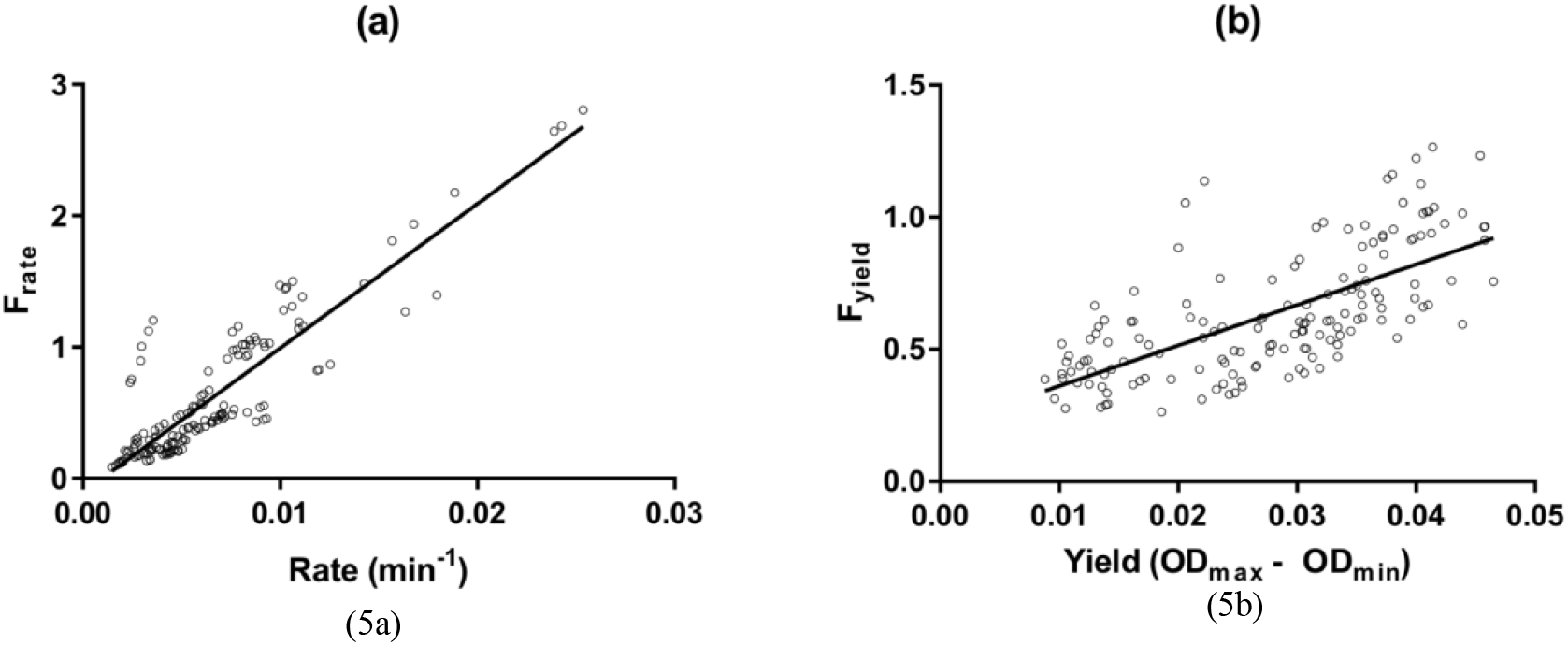
Left: plot of rate determining interaction factors vs rate with the associated regression line generated from the model: t = 23.8, df = 154, P < 0.001, multiple R^2^ = 0.79, and adjusted R^2^ = 0.79. Right: plot of the yield determining interaction factors vs rate with the associated regression line generated from the model: t = 11.5, df = 154, P < 0.001, multiple R^2^ = 0.46, and adjusted R^2^ = 0.46. Both models exclude monoculture because it is meaningless to talk about interspecies interactions in monocultures.

## 4 Discussion

We aimed to evaluate the effects of consortia diversity and species identity on the growth rates and yield of bacterial isolates growing on landfill leachate. While the majority of bioprocessing optimisation focuses on engineering single strains or manipulating process parameters, our work focused on consortia composition and rapid methods to identify consortia size and potential keystone species for growth and yield. This was under the assumption that higher growth rate indicated a faster rate of landfill leachate detoxification, and higher yield indicated a more comprehensive degradation of components within the leachate (i.e. more substrates are degraded). This assumption was due to landfill leachate providing the only carbon source available to the bacteria.

Our methodology provided small-scale, high-throughput testing of a large number of synthetic bacterial consortia combinations. The 96-well plate reader methodology allowed for automated and rapid collection of growth rate and yield data. The simplicity of the experimental design makes it robust for the testing of even higher order combinations of bacteria isolated from more complex ecosystems. It should be noted that the six bacteria tested represent only a snapshot of the full system, due to unculturable bacteria, media selection and culture conditions, which may bias the isolated strains. In this study, the procedure used for isolation of strains from landfill leachate resulted in biased isolation of aerobic heterotrophs.

Our results show that overall positive interactions resulting in either higher yields or rates are rare, implying competition could be the main interaction within our consortia. Although it might be assumptive to suggest the same for the majority of microbes within landfill leachate environments, similar findings have been revealed in other ecosystems such as those found in tree holes (Foster and Bell 2012). The only co-culture to show both positive rate and yield determining interactions was that of *Alcaligenes sp.* and *Lysinibacillus sp.,* implying an interaction possibly through the exchange of metabolites, and/or the capability to use different substrates (separate niches). In fact, *Alcaligenes sp.* is often associated with poly-aromatic hydrocarbon degradation (Kanaly and Harayama 2000), and in some cases interacts positively with other bacteria (Viñas et al. 2005). Isolates from both genera have previously been observed to efficiently reduce nitrogenous wastes from fish processing effluents, with evidence of simultaneous nitrification and denitrification (Mishra et al. 2015; Shoda and Ishikawa 2014). *Lysinibacillus sp.* were considered part of the *Bacillus* genus until differences in peptidoglycan wall composition and physiology led to its reclassification into a new genus (Ahmed et al. 2007). *Bacillus sp.* have been shown to degrade many phenolic compounds (Reddy et al. 2017) which contribute to the toxicity of typical landfill leachates. Although ubiquitous in soil, *Lysinibacillus sp.* are often associated with survival in harsh environments, demonstrating the capability to metabolise diverse environmental contaminants, for example, dichloromethanes (Wu et al. 2009) and petroleum oil hydrocarbons (Velásquez and Dussan 2009). They have also shown metal reduction capacity (Selenska-Pobell et al. 1999) where the presence of a proteinaceous S-layer on the surface of cells is thought to contribute to the biosorption of heavy metals (Velásquez and Dussan 2009). Hence it is no surprise that similar bacteria appear so often in synergistic interactions in the regression trees.

The observed predominance of competitive interactions may be due to several reasons. It may be advantageous for one microbial group to express phenotypes that equip them to compete in the presence of another microbial group, either through changes in metabolism which increase growth rate, through secretion of chemicals which slow down or stop the growth of other microbes, or even through contact killing of close proximity cells (Ghoul and Mitri 2016). Even if phenotypes are expressed that lead to increased growth rate for a particular species, they may not lead to an overall increase in growth rate of the whole consortium because the growth rates of competing species may suffer disproportionately. Moreover, if two or more microbial groups occupying separate niches detect the presence of one another, they could still express competitive phenotypes to increase their growth rate (Abisado et al. 2018) (as they would be unaware of their different respective metabolic requirements), this may be detected as an overall increase in yield i.e. the carrying capacity of the leachate would be higher as a more diverse set of compounds are utilised as substrates and degraded. However, if this were the case then one would expect to see some consortia with both *F_rate_* and *F_yieid_*> 1, which is only seen in one consortium (*Lysinibacillus sp.* and *Alcaligenes sp.*). Alternatively, these isolates could secrete chemicals into the media which hinder the growth of one another even though occupying different niches means they cannot be in direct competition. This is a “shooting oneself in the foot” situation where a consortium may have the potential to perform well but does not due to natural competition.

Another potential reason for the observed predominance of competitive interactions is that cheaters are likely to be common. Cheaters are species which consume common resources, possibly in the form of useful substrates or co-factors, to maximise their own growth without any positive contribution to the consortium. From a game theoretic view, these systems are vulnerable to exploitation (Lambert, Vyawahare, and Austin 2014). Even if there are no cheaters, the system may only be in a metastable state. It would only take one microorganism to evolve competitive phenotypes under the selective pressure of the environment to then overwhelm the other microorganisms and cause the system to crash. Hence, given enough time, even consortia that are perceived to have overall synergistic interactions may crash. This would cause issues for industrial bioprocesses, which may seek to operate continuously, as any small perturbation in the growth conditions could cause changes in the composition of the consortium ultimately leading to suboptimal performance (for this reason, it may be preferable to operate such systems in batch mode, for example, in sequential batch reactors). This may be more likely to occur when species richness increases, as more species could lead to more points of failure within the system i.e. there is a higher probability of a negative interaction existing (which ends up having a detriment on the whole system) as the number of unique species increases. Our results support this, showing that at a higher species richness rate determining and yield determining interactions become more negative.

We modelled the changes in interaction with species richness/biodiversity and found the most synergistic rate determining interactions occurred in cultures of two bacteria (co-cultures), with the ecosystem productivity reducing quickly as species richness increased. A significant correlation between species richness and rate determining interactions was seen in the form of a decaying power relationship (Figure 2a). A significant second order polynomial model was used to predict yield determining interactions with species richness, then each model was changed to a mixed model to account for community identity as a random effect. It was found that community identity is an important factor in the determination of consortia interactions. It is, however, more important in the prediction of yield determining interactions than for rate determining interactions. Simple linear regression enabled analysis of the changes of rate determining interactions with rate, and yield determining interactions with yield. As predicted, more positive rate determining interactions lead to higher rates, and the same is true for yield but to a lesser extent (Figure 5). Therefore, in order to maximise rate and yield, consortia with the most positive interactions need to be designed. No correlation was found between additive rates or yields of the species in a consortium and the actual yields or rates of the consortium (Figure 1). This indicates that simply adding the properties of monocultures to predict the properties of consortia is not a valid way of designing these ecosystems, implying that these systems are highly interactive. For instance, two very well-performing monocultures may compete very much, and two poorly performing monocultures may cooperate very much, justifying the full factorial experiment approach.

A weakness in inferring interactions by calculating the ratio of community growth rate to the sum of monoculture growth rates is that only overall interactions are predicted. For example, a value of F ≈ 1 implies that growth rates are additive, but it may be that one bacterial isolate overwhelms another, leading to a significant increase in the growth rate of the first isolate and a reduction in the other. Therefore, it cannot be said if there are both positive and negative interactions occurring within a community, only the overall interaction can be inferred. To account for this weakness, regression tree analysis was used to tease out more subtle interactions. These proved very effective in revealing previously unseen interactions without actually needing to investigate the consortia themselves with more invasive and comprehensive laboratory techniques. This provides a computational framework to infer more complex interactions from a large but basic data-set. We found a keystone species which on average resulted in an increase in consortium yields when it was present. This was *Lysinibacillus sp.* which was also found in the only consortium to have both positive yield and rate determining interactions. This implies that for breakdown of landfill leachates one should consider *Lysinibacillus sp.* in the design of synthetic microbial consortia.

Interestingly, the presence of any other bacteria into the synergistic *Alcaligenes sp.* and *Lysinibacillus sp.* co-culture had the opposite effect i.e.reducing overall rate and yield determining interactions. This may be due to initiation of a competitive phenotype when a third bacterial species enters one of their respective niches. All consortia of three, four or five bacteria that contain both of these isolates had at least one type of negative interaction (but this is true of any pair of isolates).

Regression tree analysis also revealed that, on average, the highest yield-determining interactions occurred when *Lysinibacillus sp.* was present with *Paenibacillus sp.,* implying that these species tend more often to form synergistic interactions with one another, through metabolite exchange or occupying separate niches. Regression tree analysis partitioned the yield interaction data by species presence more often than the rate interaction data (which was more often split by species richness), supporting the mixed effects models, which state that yield interactions are affected more by consortium identity than rate interactions were. The rate interaction regression tree (Figure 2) partitions the data mainly based on richness. The most positive rate interactions are seen at a low species richness. Once again *Alcaligenes sp.* appears in the regression tree but is associated with a decrease in rate-determining interactions, but only in high species richness consortia (however when combined with *Lysinibacillus sp.* it is the best overall performing co-culture in both yield and rate determining interactions). This implies that this isolate is only synergistic with a narrow range of other species, where a competitive phenotype might develop in response to bacteria with shared niche requirements.

The absence of *Stenotrophomonas chelatiphaga* (isolate 13) was associated with an average increase in both rate and yield determining interactions. This could mean *S. chelatiphaga* more often than not competes, either through cheating or expression of competitive phenotypes regardless of the environment it is put in. This would imply that while *S. chelatiphaga* tends not to compete for resources when in consortia, it does not have much of a synergistic effect either as *F_rate_* ≈ 1. This also suggests that other isolates tend to have synergistic interactions between one another in cocultures which exclude *S. chelatiphaga,* which can be seen from Figure 3 as the mean *F_rate_* is relatively high, being on average 1.37 in all co-cultures which exclude *S. chelatiphaga.*

In conclusion, we found that rate determining interspecies interactions decreased as consortium biodiversity increased, and that yield determining interspecies interactions first increased, and then decreased as biodiversity increased. Most consortia exhibit competitive traits instead of synergistic traits, with only one consortium showing positive rate and yield determining interactions. Mixed effects models indicate that consortium identity is more important in the prediction of yield determining interactions than rate determining interactions, and this was further supported by the partitioning of the data using regression tree analysis. Richness more significantly split the data than species presence for rate determining interactions, but species presence more commonly split the yield determining interaction data. We were then able to infer several specific interspecies interactions without conducting specific interspecies interaction experiments solely by virtue of having a complete (factorial) dataset of simple yield and rate data by applying the regression tree algorithm to it. This illustrates that regression tree analysis can be used as a crucial tool in speeding up the determination of interspecies interactions due to its ability to process data generated through fast high throughput techniques. From a leachate treatment perspective, the enrichment procedure would need to be expanded to capture more diverse isolates, and the same methodology described here applied to further increase yields and rates of leachate degradation. Finally, microbial diversity is often a distinct advantage in waste treatment when waste composition varies, and this would need to be tested with any prospective high performance synthetic consortia.

## Supporting information

Supplemental image of gel electrophoresis and growth curves

## Author contributions

This study was designed and coordinated by GH and JP. GH planned and undertook the practical aspects of the analyses with EK and NC. JP is the principal investigator with APB providing statistical and interpretation guidance for analysing results. The manuscript was written by GH and commented on by all authors.

## Acknowledgements

This research was funded by the EPSRC (Vacation Bursary Award), BBSRC (Impact Acceleration Award), Department of Chemical and Biological Engineering, The University of Sheffield (Summer Undergraduate Research Fellowship). JP would like to thank Marcus DuPree Thomas (Viridor Ltd.) for provision of leachate for microbial isolations. GH would like to thank all members of the EK and JP research groups for offering the space and resources to undertake this research including Adam Flinders for technical help.

## References

Abisado, Rhea G, Saida Benomar, Jennifer R Klaus, Ajai A Dandekar, and Josephine R Chandler. 2018. “Bacterial Quorum Sensing and Microbial Community Interactions.” MBio 9 (3): e02331–17. https://doi.org/10.1128/mBio.02331-17.

Ahmed, I., A. Yokota, A. Yamazoe, and T. Fujiwara. 2007. “Proposal of Lysinibacillus Boronitolerans Gen. Nov. Sp. Nov., and Transfer of Bacillus Fusiformis to Lysinibacillus Fusiformis Comb. Nov. and Bacillus Sphaericus to Lysinibacillus Sphaericus Comb. Nov.” INTERNATIONAL JOURNAL OF SYSTEMATIC AND EVOLUTIONARY MICROBIOLOGY 57 (5): 1117–25. https://doi.org/10.1099/ijs.0.63867-0.

Bates, Douglas, Martin Mächler, Ben Bolker, and Steve Walker. 2015. “Fitting Linear Mixed-Effects Models Using Lme4.” Journal of Statistical Software 67 (1): 1–48. https://doi.org/10.18637/jss.v067.i01.

Chiu, Hsuan Chao, Roie Levy, and Elhanan Borenstein. 2014. “Emergent Biosynthetic Capacity in Simple Microbial Communities.” PLoS Computational Biology 10 (7). https://doi.org/10.1371/journal.pcbi.1003695.

Fiegna, Francesca, Alejandra Moreno-Letelier, Thomas Bell, and Timothy G Barraclough. 2015. “Evolution of Species Interactions Determines Microbial Community Productivity in New Environments.” The ISME Journal 9 (5): 1235–45. https://doi.org/10.1038/ismej.2014.215.

Foster, Kevin R., and Thomas Bell. 2012. “Competition, Not Cooperation, Dominates Interactions among Culturable Microbial Species.” Current Biology 22 (19): 1845–50. https://doi.org/10.1016/j.cub.2012.08.005.

Ghoul, Melanie, and Sara Mitri. 2016. “The Ecology and Evolution of Microbial Competition.” Trends in Microbiology. https://doi.org/10.1016/j.tim.2016.06.011.

Hall, B. G., H. Acar, A. Nandipati, and M. Barlow. 2014. “Growth Rates Made Easy.” Molecular Biology and Evolution 31 (1): 232–38. https://doi.org/10.1093/molbev/mst187.

Kanaly, R A, and S Harayama. 2000. “Biodegradation of High-Molecular-Weight Polycyclic Aromatic Hydrocarbons by Bacteria.” Journal of Bacteriology 182 (8): 2059–67. http://www.ncbi.nlm.nih.gov/pubmed/10735846.

Karunakaran, E., D. Vernon, C. A. Biggs, A. Saul, D. Crawford, and H. Jensen. 2016. “Enumeration of Sulphate-Reducing Bacteria for Assessing Potential for Hydrogen Sulphide Production in Urban Drainage Systems.” Water Science and Technology 73 (12): 3087–94. https://doi.org/10.2166/wst.2016.026.

Kjeldsen, Peter, Morton A Barlaz, Alix P Rooker, Anders Baun, Anna Ledin, and Thomas H Christensen. 2002. “Present and Long-Term Composition of MSW Landfill Leachate: A Review.” Critical Reviews in Environmental Science and Technology 32 (4): 297–336. https://doi.org/10.1080/10643380290813462.

Lambert, Guillaume, Saurabh Vyawahare, and Robert H. Austin. 2014. “Bacteria and Game Theory: The Rise and Fall of Cooperation in Spatially Heterogeneous Environments.” Interface Focus 4 (4). https://doi.org/10.1098/rsfs.2014.0029.

Mishra, Saurabh S., Anoop R. Markande, Radhika P. Keluskar, Indrani Karunasagar, and Binaya B. Nayak. 2015. “Simultaneous Nitrification and Denitrification by Novel Heterotrophs in Remediation of Fish Processing Effluent.” Journal of Basic Microbiology 55 (6): 772–229. https://doi.org/10.1002/jobm.201400783.

Pandhal, Jagroop, and Josselin Noirel. 2014. “Synthetic Microbial Ecosystems for Biotechnology.” Biotechnology Letters 36 (6): 1141–51. https://doi.org/10.1007/s10529-014-1480-y.

Peralta-Yahya, Pamela P., Fuzhong Zhang, Stephen B. Del Cardayre, and Jay D. Keasling. 2012. “Microbial Engineering for the Production of Advanced Biofuels.” Nature. https://doi.org/10.1038/nature11478.

Reddy, M. Venkateswar, Yuka Yajima, Du Bok Choi, and Young Cheol Chang. 2017. “Biodegradation of Toxic Organic Compounds Using a Newly Isolated Bacillus Sp. CYR2.” Biotechnology and Bioprocess Engineering 22 (3): 339–46. https://doi.org/10.1007/s12257-017-0117-0.

Sambrook, J, and D W Russell. 2001. “Molecular Cloning: A Laboratory Manual.” Cold Spring Harbor Laboratory Press, Cold Spring Harbor, NY, 999. http://books.google.com/books?id=YTxKwWUiBeUC&printsec=frontcover%5Cnpapers2://publication/uuid/BBBF5563-6091-40C6-8B14-06ACC3392EBB.

Selenska-Pobell, Sonja, Petra Panak, Vanya Miteva, Ivo Boudakov, Gert Bernhard, and Heino Nitsche. 1999. “Selective Accumulation of Heavy Metals by Three Indigenous Bacillus Strains, B. Cereus, B. Megaterium and B. Sphaericus, from Drain Waters of a Uranium Waste Pile.” FEMSMicrobiology Ecology 29 (1): 59–67. https://doi.org/10.1016/S0168-6496(98)00130-5.

Shoda, Makoto, and Yoichi Ishikawa. 2014. “Heterotrophic Nitrification and Aerobic Denitrification of High-Strength Ammonium in Anaerobically Digested Sludge by Alcaligenes Faecalis Strain No. 4.” Journal of Bioscience and Bioengineering 117 (6): 737–41. https://doi.org/10.1016/J.JBIOSC.2013.11.018.

Simutis, Rimvydas, and Andreas Lübbert. 2015. “Bioreactor Control Improves Bioprocess Performance.” Biotechnology Journal 10 (8): 1115–30. https://doi.org/10.1002/biot.201500016.

Singh, Vineeta, Shafiul Haque, Ram Niwas, Akansha Srivastava, Mukesh Pasupuleti, and C. K.M. Tripathi. 2017. “Strategies for Fermentation Medium Optimization: An in-Depth Review.” Frontiers in Microbiology 7: 2087. https://doi.org/10.3389/fmicb.2016.02087.

Stewart, Eric J. 2012. “Growing Unculturable Bacteria.” Journal of Bacteriology. https://doi.org/10.1128/JB.00345-12.

Therneau, Terry, Beth Atkinson, Brian Ripley, and Maintainer Brian Ripley. 2015. “Rpart: Recursive Partitioning and Regression Trees.” R Package Version 4.1-10.

Velásquez, Lina, and Jenny Dussan. 2009. “Biosorption and Bioaccumulation of Heavy Metals on Dead and Living Biomass of Bacillus Sphaericus.” Journal of Hazardous Materials 167 (1-3): 713–16. https://doi.org/10.1016/j.jhazmat.2009.01.044.

Viñas, Marc, Jordi Sabaté, María José Espuny, and Anna M Solanas. 2005. “Bacterial Community Dynamics and Polycyclic Aromatic Hydrocarbon Degradation during Bioremediation of Heavily Creosote-Contaminated Soil.” Applied and Environmental Microbiology 71 (11): 7008–18. https://doi.org/10.1128/AEM.71.11.7008-7018.2005.

Wu, Shijin J., Zhihang H. Hu, Lili L. Zhang, Xiang Yu, and Jianmeng M. Chen. 2009. “A Novel Dichloromethane-Degrading Lysinibacillus Sphaericus Strain Wh22 and Its Degradative Plasmid.” Applied Microbiology and Biotechnology 82 (4): 731–40. https://doi.org/10.1007/s00253-009-1873-3.

