## Supplemental image of gel electrophoresis and growth curves for "Designing synthetic bacterial consortia for landfill leachate treatment based on community matrices and regression tree analysis"

Supplementary Material

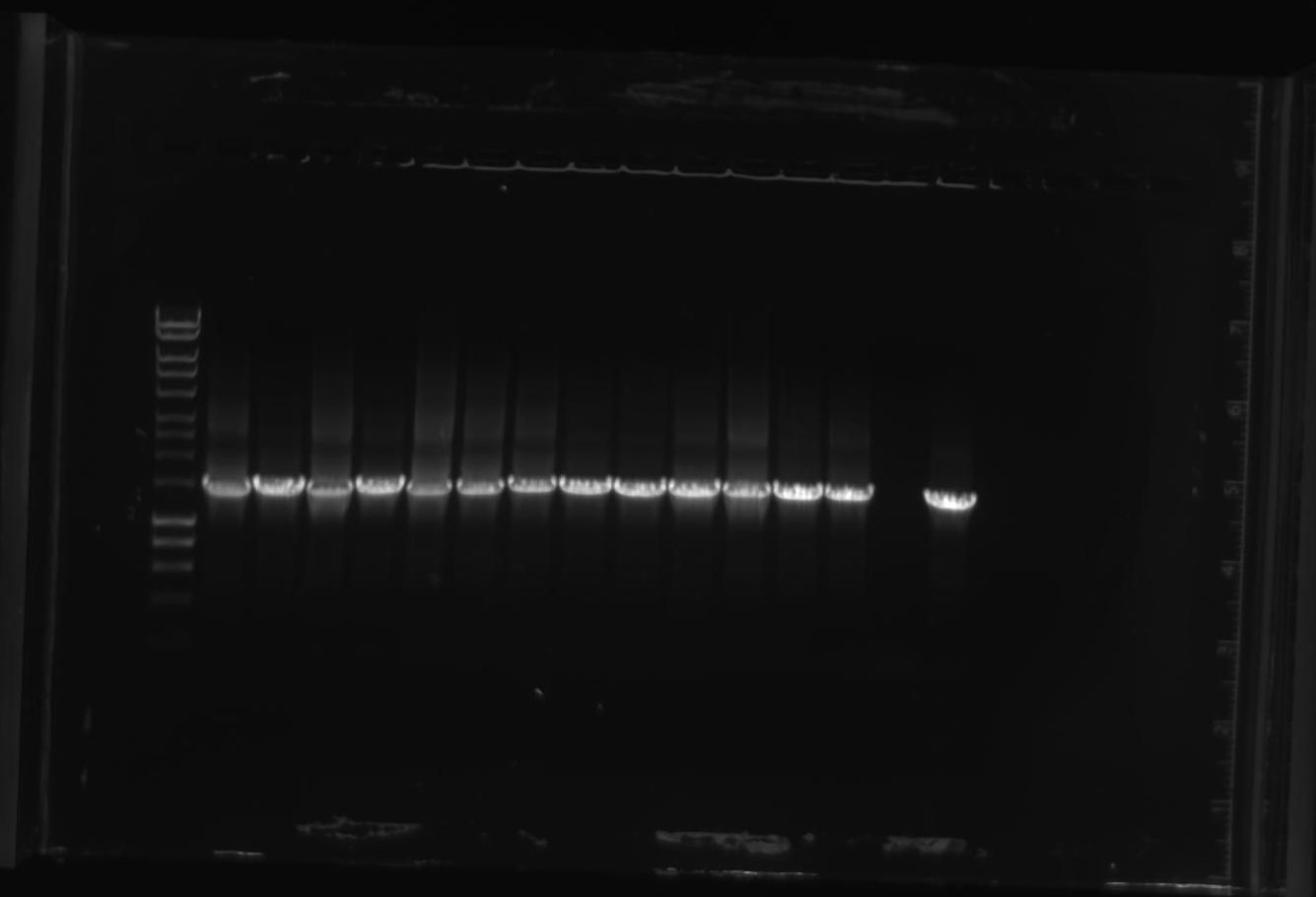

16S rDNA gel electrophoresis with Hyperladder 1KB (Bioline, UK) (left).

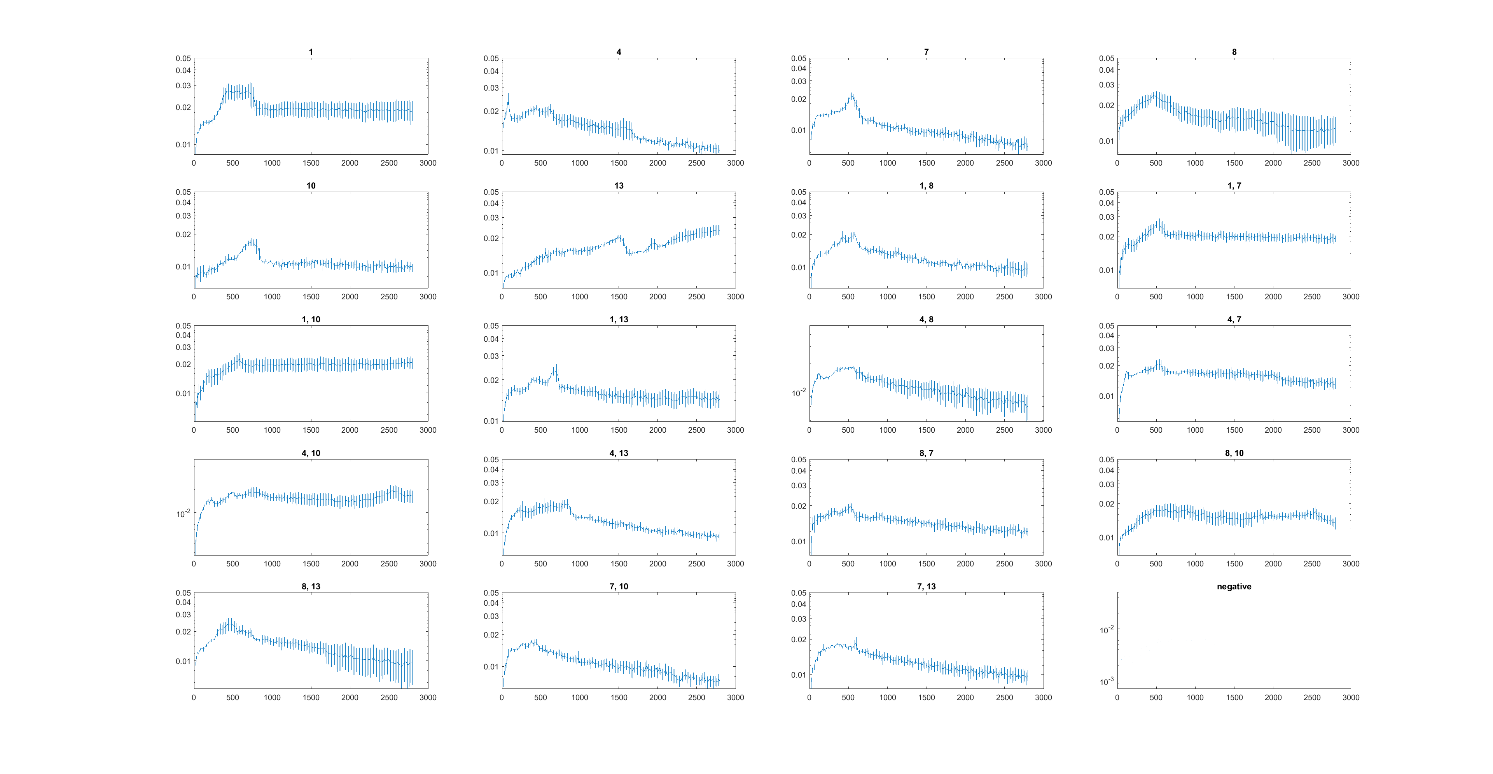

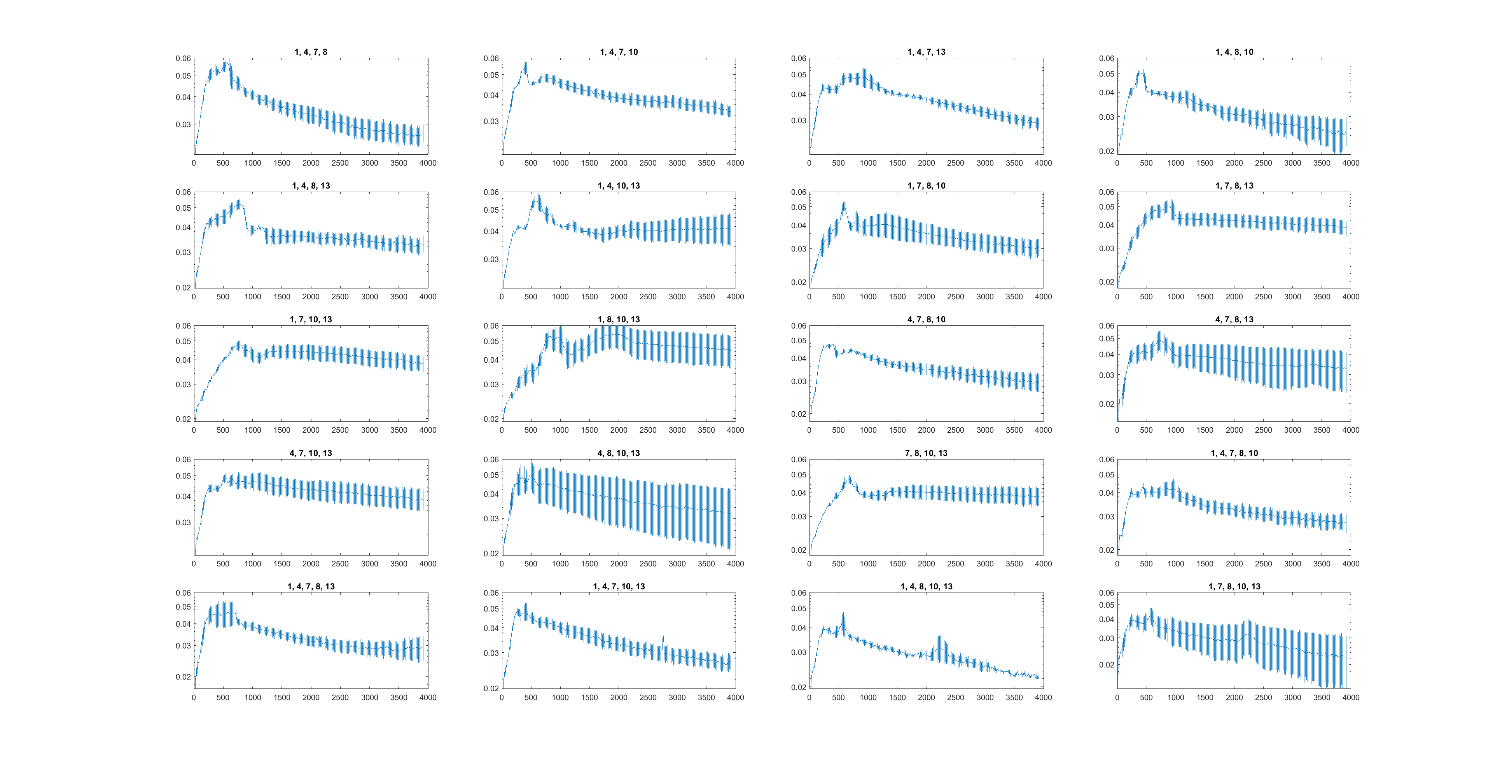

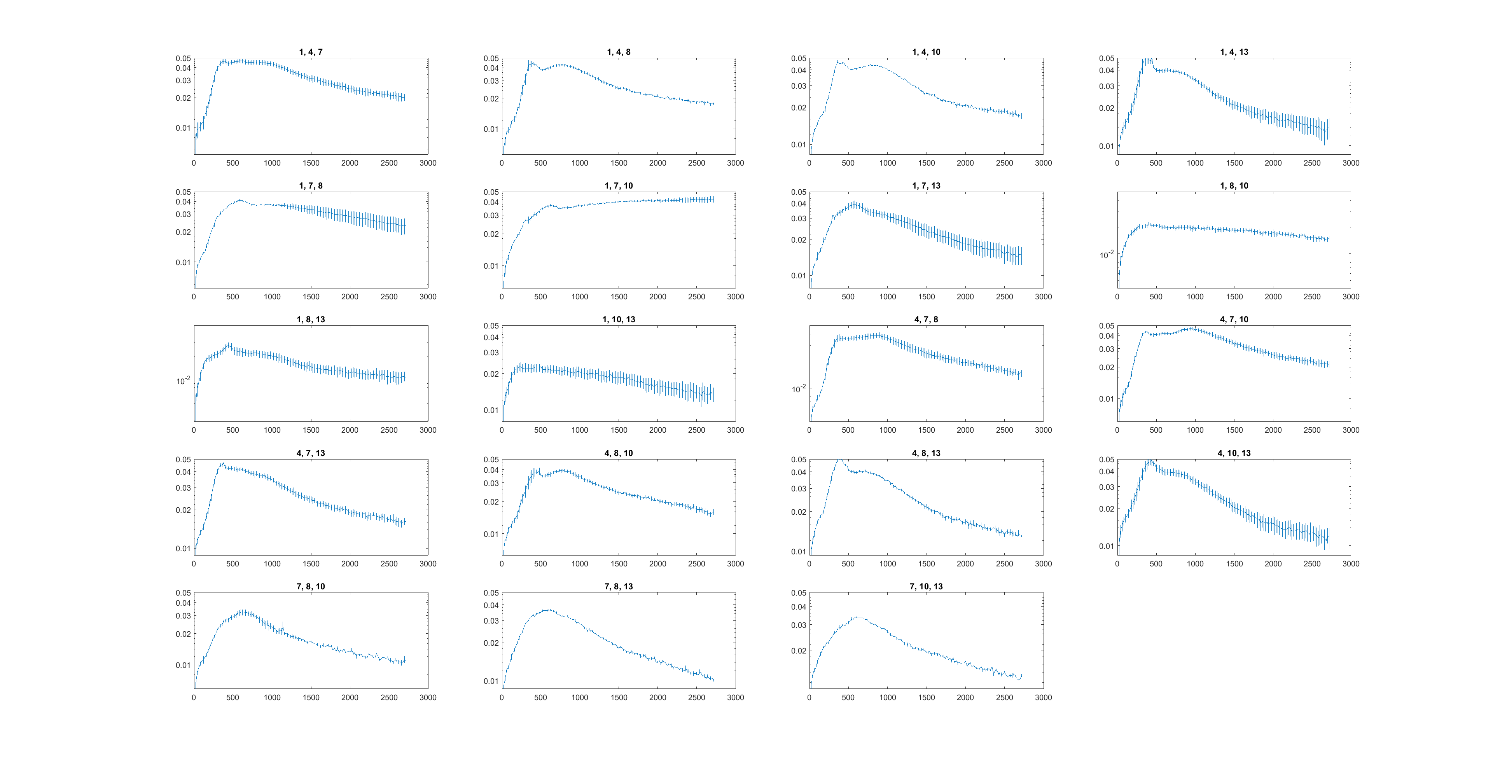

Growth measurements (x-axis: time in minutes, y-axis: optical density, error bars: SEM)
